## Supplemental Information for "Identification of putative causal loci in whole-genome sequencing data via knockoff statistics"

**He et al.**

### Supplementary Information for "Identification of putative causal loci in whole-genome sequencing data via knockoff statistics"

Zihuai He[1,2#], Linxi Liu[3], Chen Wang[4], Yann Le Guen[1], Justin Lee[2], Stephanie Gogarten[5], Fred Lu[6], Stephen Montgomery[7,8], Hua Tang[6,7], Edwin K. Silverman[9], Michael H. Cho[9], Michael Greicius[1], Iuliana Ionita-Laza[4#]

[1]Department of Neurology and Neurological Sciences, Stanford University, Stanford, CA 94305, USA
[2]Quantitative Sciences Unit, Department of Medicine, Stanford University, Stanford, CA, 94305, USA
[3]Department of Statistics, Columbia University, New York, NY 10027, USA
[4]Department of Biostatistics, Columbia University, New York, NY 10032, USA
[5]Department of Biostatistics, University of Washington, Seattle, WA, USA
[6]Department of Statistics, Stanford University, Stanford, CA, 94305, USA
[7]Department of Genetics, Stanford University, Stanford, CA, 94305, USA
[8]Department of Pathology, Stanford University, Stanford, CA, 94305, USA
[9]Channing Division of Network Medicine and Division of Pulmonary and Critical Care Medicine Division, Brigham and Women's Hospital, Harvard Medical School, Boston, MA, 02215, USA

**Supplementary Note**

**The probabilistic model for genetic variables.** As a proof of concept, we demonstrate the exchangeability of the proposed sequential knockoff generator for multivariate Gaussian distribution. The application of the proposed method to genotype dosage data is an approximation, and we demonstrate the practical performance of the proposed method by empirical studies.

The knockoff generator introduced in the paper is proposed based on a multivariate Gaussian approximation. Specifically, let $G = \left(G_1, \cdots, G_p\right)^T$ be the collection of $p$ genetic variants. We assume a multivariate normal model for $G$: $G \sim N(\mu, \Sigma)$. Based on the known haplotype block structure in the human genome, we also assume the covariance matrix $\Sigma$ is block diagonal, i.e.,

$$\Sigma = \begin{bmatrix} \Sigma_{11} & & 0 \\ & \ddots & \\ 0 & & \Sigma_{LL} \end{bmatrix},$$

where each $\Sigma_{ll}$ $(1 \leq l \leq L)$ is a $k_l$ by $k_l$ matrix. This is to say, if we divide the genome into $L$ $(L \leq p)$ contiguous non-overlapping regions/blocks, and use $\Phi_1, \cdots, \Phi_L$ to denote the set of genetic variants contained in each region respectively and let $k_l = |\Phi_l|$ $(1 \leq l \leq L)$, then with appropriately spaced and sized regions, we may use a model in which the variants from different regions are independent to each other as an approximation to the underlying correlations structure. Let $\Theta = \Sigma^{-1}$ be the precision matrix. It is easy to see that $\Theta$ is also a block diagonal matrix. For each variant $j$, let $B_j = \{j' \in [p], j' \neq j : \Theta_{jj'} \neq 0\}$. Then conditional on the variants $\{G_{j'}, j' \in B_j\}$, the variant $G_j$ is independent of the other variants. Based on our assumption, if $j \in \Phi_l$ for some $l$, then $B_j \subset \Phi_l$.

We also introduce the following notation: if $G$ is a vector of $p$ random variables, for any $A \subset [p]$, $G_A$ is defined to be the column vector $\left(G_j\right)_{j \in A}$.

When the model parameters are known, we claim that if we apply the Algorithm 1 Sequential Conditional Independent Pairs (Single Knockoff) to this model, the following claims hold at each step $j$ $(1 \leq j \leq p)$:

1. When sample $\tilde{G}_j$ from $\mathcal{L}\left(G_j \middle| G_{-j}, \tilde{G}_{1:(j-1)}\right)$ (with the convention that for $j = 1$, we sample from $\mathcal{L}\left(G_j \middle| G_{-j}\right)$), it becomes sampling $\tilde{G}_j$ from $N\left(\tilde{\mu}_j, \tilde{\sigma}_j^2\right)$, where $\tilde{\mu}_j$ is a linear combination of variants $G_{\Phi_l}$ and $\tilde{G}_{\Phi_l \cap [j-1]}$ if $j \in \Phi_l$ (for $j = 1$, it is only $G_{\Phi_l}$).
2. $\left(G, \tilde{G}_{1:j}\right)$ jointly follow a multivariate Gaussian distribution, and if we denote the precision matrix of this distribution by $\Theta^{(j+p)}$, then for any $s \in \Phi_l$, $t \in \Phi_m$ with $l \neq m$, $\Theta^{(j+p)}_{st} = 0$, $\Theta^{(j+p)}_{s,(t+p)} = 0$ when $t \leq$j, $\Theta^{(j+p)}_{(s+p),t} = 0$ when $s \leq$j, and $\Theta^{(j+p)}_{(s+p),(t+p)}$=0, when both $s \leq$j and $t \leq$j.

The claims can be shown by induction. It is easy to see that the claims hold when $j = 1$. Assume the claims hold up to step $j$. Then at step $j + 1$, as $\left(G, \tilde{G}_{1:j}\right)$ follows a multivariate normal distribution, the conditional distribution of $G_{j+1}$ given $\left(G_{-(j+1)}, \tilde{G}_{1:j}\right)$ is again a normal one. If we denote the mean of this conditional distribution as $\tilde{\mu}_{(j+1)}$, then $\tilde{\mu}_{(j+1)}$ should be a linear function of $G_{B^{(j+p)}_{j+1}}$ with $B^{(j+p)}_{j+1} = \left\{j' \neq j + 1 \colon \Theta^{(j+p)}_{j+1,j'} \neq 0, 1 \leq j' \leq p\right\}$ and $\tilde{G}_{\tilde{B}^{(j+p)}_{j+1}}$ with $\tilde{B}^{(j+p)}_{j+1} = \left\{j' - p \colon \Theta^{(j+p)}_{j+1,j'} \neq 0, p < j' \leq p + j\right\}$. Based on the second induction hypothesis at step $j$, if $j + 1 \in \Phi_l$ for some $l$, then $G_{B^{(j+p)}_{j+1}} \subset \Phi_l$ and $\tilde{G}_{\tilde{B}^{(j+p)}_{j+1}} \subset \Phi_l \cap [j]$. Therefore, the first induction hypothesis still holds at step $j + 1$.

To show the second part, without loss of generality, we can switch the order of the variables to make $G_{j+1}$ the last variable, and the corresponding precision matrix is denoted as

$$\overline{\Theta}^{(j+p)} = \begin{bmatrix} \overline{\Theta}_1^{(j+p)} & \bar{\theta}_{j+1} \\ \bar{\theta}_{j+1}^T & \Theta_{j+1,j+1}^{(j+p)} \end{bmatrix},$$

where $\overline{\Theta}_1^{(j+p)}$ is a $p+j-1$ by $p+j-1$ matrix obtained by removing the $(j+1)$st column and $(j+1)$st row from $\Theta^{(j+p)}$, $\bar{\theta}_{j+1}$ is a column vector of length $p+j-1$, obtained by removing the $(j+1)$st element from the $(j+1)$st column of $\Theta^{(j+p)}$, and $\Theta_{j+1,j+1}^{(j+p)}$ is the $(j+1)$st diagonal element of $\Theta^{(j+p)}$. Then after we sample $\tilde{G}_{j+1}$ independently from $\mathcal{L}\left(G_{j+1} \middle| G_{-(j+1)}, \tilde{G}_{1:j}\right)$, the joint distribution of $\left(G_{-(j+1)}, \tilde{G}_{1:j}, G_{j+1}, \tilde{G}_{j+1}\right)$ is still a multivariate normal one, and its precision matrix is

$$\overline{\Theta}^{(j+p+1)} = \begin{bmatrix} \overline{\Theta}_1^{(j+p)} + \left(\Theta_{j+1,j+1}^{(j+p)}\right)^{-1} \bar{\theta}_{j+1}\bar{\theta}_{j+1}^T & \bar{\theta}_{j+1} & \bar{\theta}_{j+1} \\ \bar{\theta}_{j+1}^T & \Theta_{j+1,j+1}^{(j+p)} & 0 \\ \bar{\theta}_{j+1}^T & 0 & \Theta_{j+1,j+1}^{(j+p)} \end{bmatrix}.$$

Based on this, after rearrange the order of variables, we can get the precision matrix of the joint distribution of $\left(G, \tilde{G}_{1:(j+1)}\right)$, which is denoted as $\Theta^{(j+p+1)}$. Based on the second induction hypothesis, we still have for any $s \in \Phi_l$, $t \in \Phi_m$ with $l \neq m$, $\Theta_{st}^{(j+p+1)} = 0$, $\Theta_{s,(t+p)}^{(j+p+1)} = 0$ when $t \leq j+1$, $\Theta_{(s+p),t}^{(j+p+1)} = 0$ when $s \leq j+1$, and $\Theta_{(s+p),(t+p)}^{(j+p+1)}=0$, when both $s \leq j+1$ and $t \leq j+1$. This finishes the proof for the two claims. A similar argument can also be applied to the multiple knockoffs. That is, if a same model is imposed on the original variables, then when applying Algorithm 2 to generate knockoffs, the conditional distribution is a normal one, and conditional on the nearby variants and their already constructed knockoffs, the $j$th variable is independent of the other original and knockoff variables.

In practice, as the model parameters are unknown, we use the following methods to estimate the model parameters, and approximately sample from the conditional distribution $\mathcal{L}\left(G_j \middle| G_{-j}, \tilde{G}_{1:(j-1)}\right)$ at each step:

1. Estimate the conditional mean by running a regression. Based on the first claim, the conditional mean should be a linear function of the nearby variants (those that belong to the same LD block) and their knockoffs, so we can estimate the conditional mean by regressing $G_j$ on those variables. In this paper, we choose to include the variants in a nearby region (within 200kb), under the assumption that such a region is large enough to cover an LD block.

2. We did not estimate the conditional variance, instead, we permute the residuals as an approximation to sampling from the conditional distribution.

It is worthy to note that when the size of the blocks is large enough, we may think that such a model is a reasonable approximation to the true correlation structure of genetic variants. However, the larger the block, the higher the computational cost. To make a trade-off between the computational cost and model accuracy, in this paper we set the size of the block to be about 200kb (+/-100kb from the target variant), as the typical LD block is less than 100kb. Within these blocks LD decreases slowly with physical distance, but between blocks LD decays rapidly[63].

**Proof of the exchangeability property of the sequential model for multiple knockoffs.** Let $G = (G_1, \cdots, G_p)$ be $p$ original explanatory variables (i.e. genetic variants in our case), also denoted as $\tilde{G}^0$; $\tilde{G}^{\mathrm{m}} = (\tilde{G}_1^m, \cdots, \tilde{G}_p^m)$, $1 \le m \le M$, be $M$ ($M \ge 2$) groups of knockoff features. $\sigma = (\sigma_j)_{1 \le j \le p}$ is a collection of $p$ permutations over the set of integers $\{0, 1, \cdots, M\}$, with each $\sigma_j$ corresponding to an original feature $X_j$. The variables after swapping are defined as $(\tilde{G}^0, \tilde{G}^1, \cdots, \tilde{G}^{\mathrm{M}})_{\mathrm{swap}(\sigma)} := (U^0, U^1, \cdots, U^{\mathrm{M}})$, with $U_j^{\mathrm{m}} = \tilde{G}_j^{\sigma_j(m)}$ for all $1 \le j \le p, 1 \le m \le M$. The extended exchangeability condition for multiple knockoffs is defined as follows.

*Definition 1. The multiple knockoffs* $(\tilde{G}^0, \tilde{G}^1, \cdots, \tilde{G}^M)$ *satisfy the extended exchangeability condition if* $\left(\tilde{G}^0, \tilde{G}^1, \cdots, G^M\right)_{swap(\sigma)}$ *follows the same distribution as* $\left(\tilde{G}^0, \tilde{G}^1, \cdots, \tilde{G}^M\right)$ *for any* $\sigma$.

We prove that if we generate multiple knockoffs by applying Algorithm 2, the extended exchangeability is satisfied. We denote the probability mass function (PMF) of $(G_{1:p}, \tilde{G}_{1:j-1}^1, \cdots, \tilde{G}_{1:j-1}^{\mathrm{M}})$ as $\mathcal{L}\left(G_{-j}, G_j, \tilde{G}_{1:j-1}^1, \cdots, \tilde{G}_{1:j-1}^{\mathrm{M}}\right)$. Our argument is based on induction with the following induction hypothesis: after $j$ steps, for all $1 \le l \le j$, the variables $\left(\tilde{G}_l^0, \tilde{G}_l^1, \cdots, \tilde{G}_l^M\right)$ are exchangeable with respect to any permutation $\sigma_l$ over the set of integers $\{0, 1, \cdots, M\}$ in the joint distribution $\mathcal{L}\left(G_{1:p}, \tilde{G}_{1:j}^1, \cdots, \tilde{G}_{1:j}^M\right)$.

It is easy to check the induction hypothesis is true when $j = 1$. Next, assuming the induction hypothesis holds for the first $j - 1$, we show that it also holds after $j$ steps. At step $j$, $\tilde{G}_j^1, \cdots, \tilde{G}_j^{\mathrm{M}}$ are conditionally independent and follow the same distribution. The conditional PMF of $\tilde{G}_j^{(1)}$ given $G_{1:p}, \tilde{G}_{1:j-1}^1, \cdots, \tilde{G}_{1:j-1}^{\mathrm{M}}$ is

$$\frac{\mathcal{L}\left(G_{-j}, \tilde{G}_j^1, \tilde{G}_{1:j-1}^1, \cdots, \tilde{G}_{1:j-1}^{\mathrm{M}}\right)}{\sum_u \mathcal{L}\left(G_{-j}, u, \tilde{G}_{1:j-1}^1, \cdots, \tilde{G}_{1:j-1}^{\mathrm{M}}\right)}.$$

Then the joint PMF of $\left(G_{1:p}, \tilde{G}_{1:j}^1, \cdots, \tilde{G}_{1:j}^{\mathrm{M}}\right)$ is the product of the conditional PMF with the joint PMF of $\left(G_{1:p}, \tilde{G}_{1:j-1}^1, \cdots, \tilde{G}_{1:j-1}^{\mathrm{M}}\right)$:

$$\frac{\prod_{m=0}^{M} \mathcal{L}\left(G_{-j}, \tilde{G}_j^{\mathrm{m}}, \tilde{G}_{1:j-1}^1, \cdots, \tilde{G}_{1:j-1}^{\mathrm{M}}\right)}{\left(\sum_u \mathcal{L}\left(G_{-j}, u, \tilde{G}_{1:j-1}^1, \cdots, \tilde{G}_{1:j-1}^{\mathrm{M}}\right)\right)^M}.$$

The PMF remains invariant with respect to any permutation of $\left(\tilde{G}_j^0, \tilde{G}_j^1, \cdots, \tilde{G}_j^{\mathrm{M}}\right)$. Based on the induction hypothesis, we also have for any $l < j$, the joint distribution $\mathcal{L}$ is symmetric in $\left(\tilde{G}_l^0, \tilde{G}_l^1, \cdots, \tilde{G}_l^{\mathrm{M}}\right)$. Combining these two facts, the induction hypothesis holds after $j$ steps.

**Proof of FDR control.** For *KnockoffScreen*, we construct multiple groups of knockoff features and introduce a new feature importance statistic $\tau_{\Phi_{kl}} = T_{\Phi_{kl}}^{(0)} - \underset{1 \le m \le M}{\text{median}}\, T_{\Phi_{kl}}^{(m)}$ instead of $T_{\Phi_{kl}}^{(0)} - T_{\Phi_{kl}}^{(1)}$, where $\Phi_{kl}$ denotes a window on the genome. In this section, we show what with the newly introduced feature important statistic, the method still leads to valid FDR control.

Let $\Phi_{k_1 l_1}, \Phi_{k_2 l_2}, \cdots, \Phi_{k_W l_W}$ be a set of non-overlapping windows on the genome. Recall that each window $\Phi_{k_\omega l_\omega}$ $(1 \le \omega \le W)$ is defined to be $\Phi_{k_\omega l_\omega} = \{j: k_\omega \le j \le l_\omega\} \subset [p]$. In other words, $\Phi_{k_\omega l_\omega}$'s are disjoint subsets of $[p]$. $\mathcal{H}_0 = \{j: G_j \textit{ is a noncausal variant}\}$. For each window, we have introduced a pair of test statistics: $\tau_{\Phi_{k_\omega l_\omega}}$ and $\kappa_{\Phi_{k_\omega l_\omega}}$. Similar to all types of knockoff filters, the FDR control is achieved based on the following key property of the test statistics.

*Property 1*: conditioning on $\left(\tau_{\Phi_{k_\omega l_\omega}}\right)_{1 \le \omega \le W}$ and $\kappa_{\Phi_{k_\omega l_\omega}}$'s for non-null windows, $\kappa_{\Phi_{k_\omega l_\omega}}$ for null windows are i.i.d. random variables uniformly distributed on $\{0, 1, \cdots, M\}$.

Under multiple knockoffs framework, the property can be view as an extension of the sign-flipping one corresponding to the single knockoffs.

To show this property, we consider a collection of permutations $\left(\sigma_{\Phi_{k_\omega l_\omega}}\right)_{1 \le \omega \le W}$ on $\{0, 1, \cdots, M\}$ defined in the following way: if $\Phi_{k_\omega l_\omega} \subset \mathcal{H}_0$ , then $\sigma_{\Phi_{k_\omega l_\omega}}$ can be any permutation; otherwise, $\sigma_{\Phi_{k_\omega l_\omega}}$ is the identity permutation. We will show the following two sets of random variables follow the same distribution:

$$\left(\left(\sigma_{\Phi_{k_\omega l_\omega}}\left(\kappa_{\Phi_{k_\omega l_\omega}}\right)\right)_{1 \le \omega \le W}, \left(\tau_{\Phi_{k_\omega l_\omega}}\right)_{1 \le \omega \le W}\right) \sim \left(\left(\kappa_{\Phi_{k_\omega l_\omega}}\right)_{1 \le \omega \le W}, \left(\tau_{\Phi_{k_\omega l_\omega}}\right)_{1 \le \omega \le W}\right)$$

The proof is based on the following observation: for window $\Phi_{k_\omega l_\omega}$ with test statistics $\tau_{\Phi_{k_\omega l_\omega}}$ and $\kappa_{\Phi_{k_\omega l_\omega}}$, if we apply any permutation $\sigma_{\Phi_{k_\omega l_\omega}}$ to variables and their knockoffs corresponding to the variants covered by the same window, then based on the permuted data set the test statistics are exactly $\tau_{\Phi_{k_\omega l_\omega}}$ and $\sigma_{\Phi_{k_\omega l_\omega}}\left(\kappa_{\Phi_{k_\omega l_\omega}}\right)$. More precisely, if we define $(\widetilde{\boldsymbol{G}}^{\mathbf{0}}, \widetilde{\boldsymbol{G}}^{\mathbf{1}}, \cdots, \widetilde{\boldsymbol{G}}^{\mathbf{M}})_{\text{swap}} := (\boldsymbol{U}^{\mathbf{0}}, \boldsymbol{U}^{\mathbf{1}}, \cdots, \boldsymbol{U}^{\mathbf{M}})$ as $\boldsymbol{U}_{\boldsymbol{j}}^{\mathbf{m}} = \widetilde{\boldsymbol{G}}_{\boldsymbol{j}}^{\sigma_{\Phi_{k_\omega l_\omega}}(\boldsymbol{m})}$ for $j \in \Phi_{k_\omega l_\omega}$, and denote the test statistics for window $\Phi_{k_\omega l_\omega}$ based on $(\widetilde{\boldsymbol{G}}^{\mathbf{0}}, \widetilde{\boldsymbol{G}}^{\mathbf{1}}, \cdots, \widetilde{\boldsymbol{G}}^{\mathbf{M}})_{\text{swap}}$ as $\hat{\sigma}_{\Phi_{k_\omega l_\omega}}$ and $\hat{\tau}_{\Phi_{k_\omega l_\omega}}$, then $\hat{\tau}_{\Phi_{k_\omega l_\omega}} = \tau_{\Phi_{k_\omega l_\omega}}$ and $\hat{\sigma}_{\Phi_{k_\omega l_\omega}} = \sigma_{\Phi_{k_\omega l_\omega}}\left(\kappa_{\Phi_{k_\omega l_\omega}}\right)$. In combination with the fact that $(\widetilde{\boldsymbol{G}}^{\mathbf{0}}, \widetilde{\boldsymbol{G}}^{\mathbf{1}}, \cdots, \widetilde{\boldsymbol{G}}^{\mathbf{M}})_{\text{swap}}$ and $(\widetilde{\boldsymbol{G}}^{\mathbf{0}}, \widetilde{\boldsymbol{G}}^{\mathbf{1}}, \cdots, \widetilde{\boldsymbol{G}}^{\boldsymbol{M}})$ follow the same distribution, we have

$$\left(\left(\sigma_{\Phi_{k_\omega l_\omega}}\left(\kappa_{\Phi_{k_\omega l_\omega}}\right)\right)_{1 \le \omega \le W}, \left(\tau_{\Phi_{k_\omega l_\omega}}\right)_{1 \le \omega \le W}\right)$$

$$= f\left(\left(\widetilde{\boldsymbol{G}}^{\mathbf{0}}, \widetilde{\boldsymbol{G}}^{\mathbf{1}}, \cdots, \widetilde{\boldsymbol{G}}^{\mathbf{M}}\right)_{\text{swap}}, \boldsymbol{Y}\right)$$

$$\sim f\left(\left(\widetilde{\boldsymbol{G}}^{\mathbf{0}}, \widetilde{\boldsymbol{G}}^{\mathbf{1}}, \cdots, \widetilde{\boldsymbol{G}}^{\boldsymbol{M}}\right), \boldsymbol{Y}\right)$$

$$= \left(\left(\kappa_{\Phi_{k_\omega l_\omega}}\right)_{1 \le \omega \le W}, \left(\tau_{\Phi_{k_\omega l_\omega}}\right)_{1 \le \omega \le W}\right)$$

Here $f$ is a function describing how we calculate test statistics based on the data. This finishes the proof for *Property 1.*

If *Property 1* holds, the knockoff filter is a special case of the Second Sequential Testing Procedure discussed by Barber and Candès [13], the FDR control can be obtained by using a similar argument as that used by Gimenez and Zou [30].

#### Supplementary Tables

**Table S1: Empirical evaluation of *KnockoffScreen* in the presence of population stratification driven by rare variants.** Each cell presents the empirical FDR. $\gamma$ quantifies the magnitude of population stratification; C: continuous trait; D: dichotomous trait. KnockoffScreen controls FDR at 0.10; Association Testing is based on the usual Bonferroni correction (0.05/number of tests), controlling FWER at 0.05 level.

| $\gamma$ | Trait | KnockoffScreen | KnockoffScreen 10 PCs | Association Testing | Association Testing 10 PCs |
|---|---|---|---|---|---|
| 0 | C | 0.098 | 0.102 | 0.022 | 0.020 |
| 0.25 | C | 0.084 | 0.096 | 0.094 | 0.024 |
| 0.5 | C | 0.124 | 0.084 | 0.430 | 0.018 |
| 0.75 | C | 0.196 | 0.068 | 0.926 | 0.028 |
| 0 | D | 0.106 | 0.112 | 0.056 | 0.058 |
| 0.25 | D | 0.108 | 0.100 | 0.184 | 0.042 |
| 0.5 | D | 0.198 | 0.110 | 0.846 | 0.030 |
| 0.75 | D | 0.312 | 0.090 | 0.996 | 0.034 |

**Table S2 Tissue grouping of GenoNet scores.** The GenoNet scores were trained using epigenetic annotations from the Roadmap Epigenomics Project across 127 tissues/cell types.

| Epigenome ID (EID) | Standardized Epigenome name | Group |
|---|---|---|
| E062 | Primary mononuclear cells fromperipheralblood | Blood |
| E034 | Primary T cells fromperipheralblood | Blood |
| E045 | Primary T cells e_ector/memory enriched from peripheral blood | Blood |
| E033 | Primary T cells from cord blood | Blood |
| E044 | Primary T regulatory cells fromperipheralblood | Blood |
| E043 | Primary T helper cells fromperipheralblood | Blood |
| E039 | Primary T helper naive cells fromperipheralblood | Blood |
| E041 | Primary T helper cells PMA-I stimulated | Blood |
| E042 | Primary T helper 17 cells PMA-I stimulated | Blood |
| E040 | Primary T helper memory cells from peripheral blood 1 | Blood |
| E037 | Primary T helper memory cells from peripheral blood 2 | Blood |
| E048 | Primary T CD8+ memory cells from peripheral blood | Blood |
| E038 | Primary T helper naive cells from peripheral blood | Blood |
| E047 | Primary T CD8+ naive cells from peripheral blood | Blood |
| E029 | Primary monocytes from peripheral blood | Blood |
| E031 | Primary B cells from cord blood | Blood |
| E035 | Primary hematopoietic stem cells | Blood |
| E051 | Primary hematopoietic stem cells G-CSF-mobilized Male | Blood |
| E050 | Primary hematopoietic stem cells G-CSF-mobilized Female | Blood |
| E036 | Primary hematopoietic stem cells short term culture | Blood |
| E032 | Primary B cells from peripheral blood | Blood |

| | | |
|---|---|---|
| E046 | Primary Natural Killer cells from peripheral blood | Blood |
| E030 | Primary neutrophils from peripheral blood | Blood |
| E112 | Thymus | Blood |
| E093 | Fetal Thymus | Blood |
| E115 | Dnd41 TCell Leukemia Cell Line | Blood |
| E116 | GM12878 Lymphoblastoid Cells | Blood |
| E123 | K562 Leukemia Cells | Blood |
| E124 | Monocytes-CD14+ RO01746 Primary Cells | Blood |
| E071 | Brain Hippocampus Middle | Brain |
| E074 | Brain Substantia Nigra | Brain |
| E068 | Brain Anterior Caudate | Brain |
| E069 | Brain Cingulate Gyrus | Brain |
| E072 | Brain Inferior Temporal Lobe | Brain |
| E067 | Brain Angular Gyrus | Brain |
| E073 | Brain Dorsolateral Prefrontal Cortex | Brain |
| E017 | MR90 fetal lung _broblasts Cell Line | ConnectiveTissue |
| E026 | Bone Marrow Derived Cultured Mesenchymal Stem Cells | ConnectiveTissue |
| E049 | Mesenchymal Stem Cell Derived Chondrocyte Cultured Cells | ConnectiveTissue |
| E025 | Adipose Derived Mesenchymal Stem Cell Cultured Cells | ConnectiveTissue |
| E023 | Mesenchymal Stem Cell Derived Adipocyte Cultured Cells | ConnectiveTissue |
| E052 | Muscle Satellite Cultured Cells | ConnectiveTissue |
| E055 | Foreskin Fibroblast Primary Cells skin01 | ConnectiveTissue |
| E056 | Foreskin Fibroblast Primary Cells skin02 | ConnectiveTissue |
| E057 | Foreskin Keratinocyte Primary Cells skin02 | ConnectiveTissue |
| E058 | Foreskin Keratinocyte Primary Cells skin03 | ConnectiveTissue |
| E028 | Breast variant Human Mammary Epithelial Cells (vHMEC) | ConnectiveTissue |
| E114 | A549 EtOH 0.02pct Lung Carcinoma Cell Line | ConnectiveTissue |
| E117 | HeLa-S3 Cervical Carcinoma Cell Line | ConnectiveTissue |
| E119 | HMEC Mammary Epithelial Primary Cells | ConnectiveTissue |
| E120 | HSMM Skeletal Muscle Myoblasts Cells | ConnectiveTissue |
| E121 | HSMM cell derived Skeletal Muscle Myotubes Cells | ConnectiveTissue |
| E122 | HUVEC Umbilical Vein Endothelial Primary Cells | ConnectiveTissue |
| E125 | NH-A Astrocytes Primary Cells | ConnectiveTissue |
| E126 | NHDF-Ad Adult Dermal Fibroblast Primary Cells | ConnectiveTissue |
| E127 | NHEK-Epidermal Keratinocyte Primary Cells | ConnectiveTissue |
| E128 | NHLF Lung Fibroblast Primary Cells | ConnectiveTissue |
| E129 | Osteoblast Primary Cells | ConnectiveTissue |
| E054 | Ganglion Eminence derived primary cultured neurospheres | FetalBrain |

| | | |
|---|---|---|
| E053 | Cortex derived primary cultured neurospheres | FetalBrain |
| E070 | Brain Germinal Matrix | FetalBrain |
| E082 | Fetal Brain Female | FetalBrain |
| E081 | Fetal Brain Male | FetalBrain |
| E013 | hESC Derived CD56+ Mesoderm Cultured Cells | FetalTissue1 |
| E005 | H1 BMP4 Derived Trophoblast Cultured Cells | FetalTissue1 |
| E006 | H1 Derived Mesenchymal Stem Cells | FetalTissue1 |
| E083 | Fetal Heart | FetalTissue1 |
| E099 | Placenta Amnion | FetalTissue1 |
| E089 | Fetal Muscle Trunk | FetalTissue2 |
| E090 | Fetal Muscle Leg | FetalTissue2 |
| E092 | Fetal Stomach | FetalTissue2 |
| E088 | Fetal Lung | FetalTissue2 |
| E080 | Fetal Adrenal Gland | FetalTissue2 |
| E091 | Placenta | FetalTissue2 |
| E085 | Fetal Intestine Small | Gastrointestinal |
| E084 | Fetal Intestine Large | Gastrointestinal |
| E109 | Small Intestine | Gastrointestinal |
| E106 | Sigmoid Colon | Gastrointestinal |
| E075 | Colonic Mucosa | Gastrointestinal |
| E101 | Rectal Mucosa Donor 29 | Gastrointestinal |
| E102 | Rectal Mucosa Donor 31 | Gastrointestinal |
| E110 | Stomach Mucosa | Gastrointestinal |
| E077 | Duodenum Mucosa | Gastrointestinal |
| E066 | Liver | Gastrointestinal |
| E118 | HepG2 Hepatocellular Carcinoma Cell Line | Gastrointestinal |
| E059 | Foreskin Melanocyte Primary Cells skin01 | InternalOrgans |
| E061 | Foreskin Melanocyte Primary Cells skin03 | InternalOrgans |
| E027 | Breast Myoepithelial Primary Cells | InternalOrgans |
| E100 | Psoas Muscle | InternalOrgans |
| E104 | Right Atrium | InternalOrgans |
| E095 | Left Ventricle | InternalOrgans |
| E105 | Right Ventricle | InternalOrgans |
| E065 | Aorta | InternalOrgans |
| E079 | Esophagus | InternalOrgans |
| E094 | Gastric | InternalOrgans |
| E086 | Fetal Kidney | InternalOrgans |
| E097 | Ovary | InternalOrgans |

| | | |
|---|---|---|
| E087 | Pancreatic Islets | InternalOrgans |
| E098 | Pancreas | InternalOrgans |
| E096 | Lung | InternalOrgans |
| E113 | Spleen | InternalOrgans |
| E063 | Adipose Nuclei | Muscle |
| E108 | Skeletal Muscle Female | Muscle |
| E107 | Skeletal Muscle Male | Muscle |
| E078 | Duodenum Smooth Muscle | Muscle |
| E076 | Colon Smooth Muscle | Muscle |
| E103 | Rectal Smooth Muscle | Muscle |
| E111 | Stomach Smooth Muscle | Muscle |
| E002 | ES-WA7 Cells | StemCell |
| E008 | H9 Cells | StemCell |
| E001 | ES-I3 Cells | StemCell |
| E015 | HUES6 Cells | StemCell |
| E014 | HUES48 Cells | StemCell |
| E016 | HUES64 Cells | StemCell |
| E003 | H1 Cells | StemCell |
| E024 | ES-UCSF4 Cells | StemCell |
| E020 | iPS-20b Cells | StemCell |
| E019 | iPS-18 Cells | StemCell |
| E018 | iPS-15b Cells | StemCell |
| E021 | iPS DF 6.9 Cells | StemCell |
| E022 | iPS DF 19.11 Cells | StemCell |
| E007 | H1 Derived Neuronal Progenitor Cultured Cells | StemCell |
| E009 | H9 Derived Neuronal Progenitor Cultured Cells | StemCell |
| E010 | H9 Derived Neuron Cultured Cells | StemCell |
| E012 | hESC Derived CD56+ Ectoderm Cultured Cells | StemCell |
| E011 | hESC Derived CD184+ Endoderm Cultured Cells | StemCell |
| E004 | H1 BMP4 Derived Mesendoderm Cultured Cells | StemCell |

### Supplementary Figures

**Figure S1 Distribution of power and false discovery proportion (FDP) at target FDR level 0.1 in simulation studies.** The results are based on 1000 replicates and the same settings as in Figure 1.

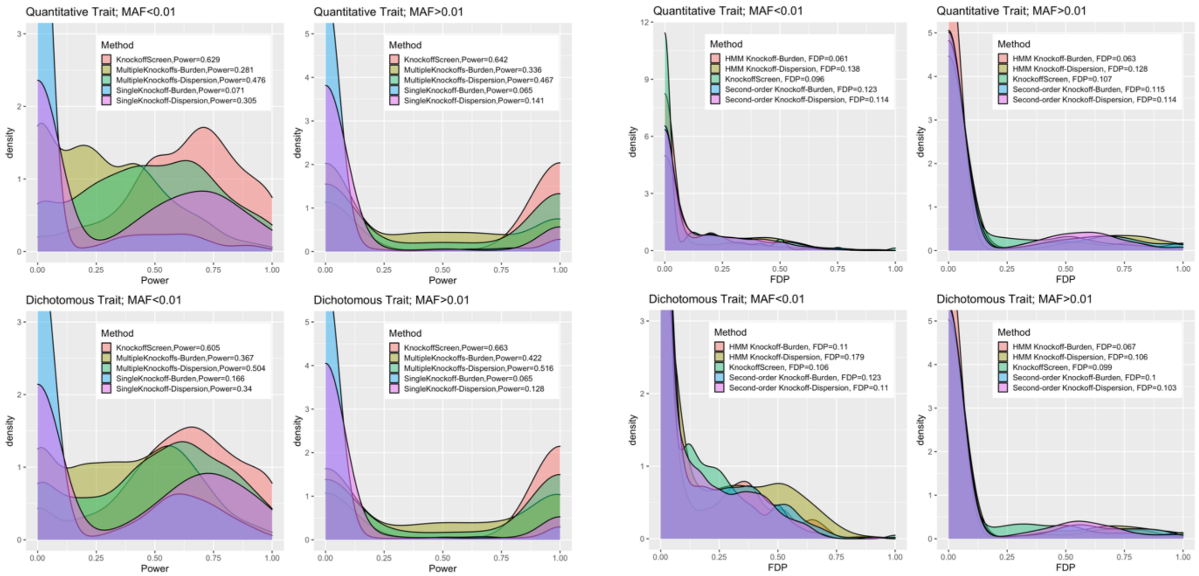

**Figure S2: Empirical validation of the extended exchangeability for rare variants.** We generated 10,000 individuals with genetic data for a 200 kb region containing 1000 genetic variants, simulated using a coalescent model (COSI). To validate the extended exchangeability, we generated two knockoffs using the proposed algorithm and evaluated whether the second order (covariance between each pair of genetic variants) is exchangeable for both rare and common variants in the regions.

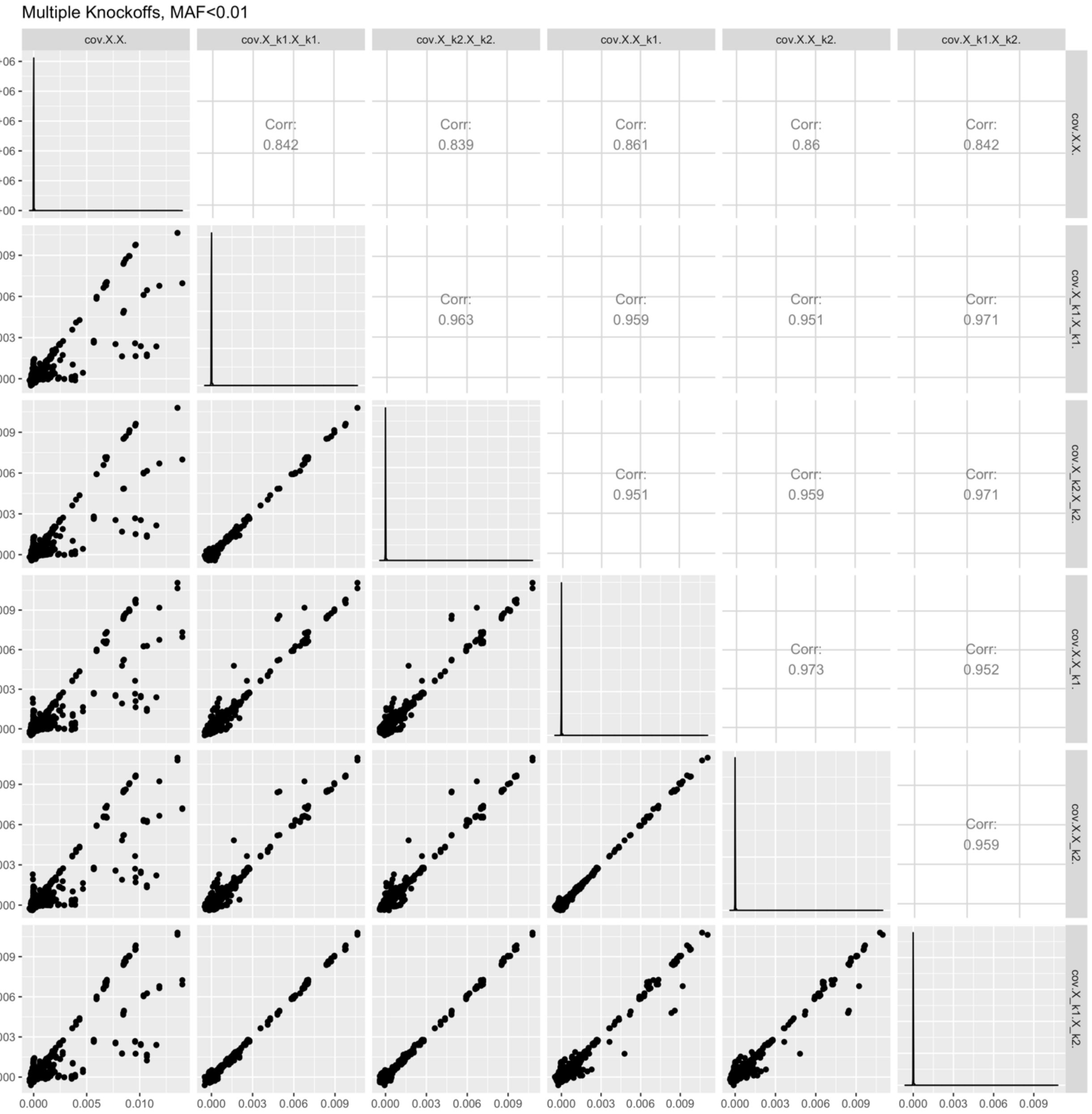

**Figure S3: Empirical validation of the extended exchangeability for common variants.** We generated 10,000 individuals with genetic data for a 200 kb region containing 1000 genetic variants, simulated using a coalescent model (COSI). To validate the extended exchangeability, we generated two knockoffs using the proposed algorithm and evaluated whether the second order (covariance between each pair of genetic variants) is exchangeable for both rare and common variants in the regions.

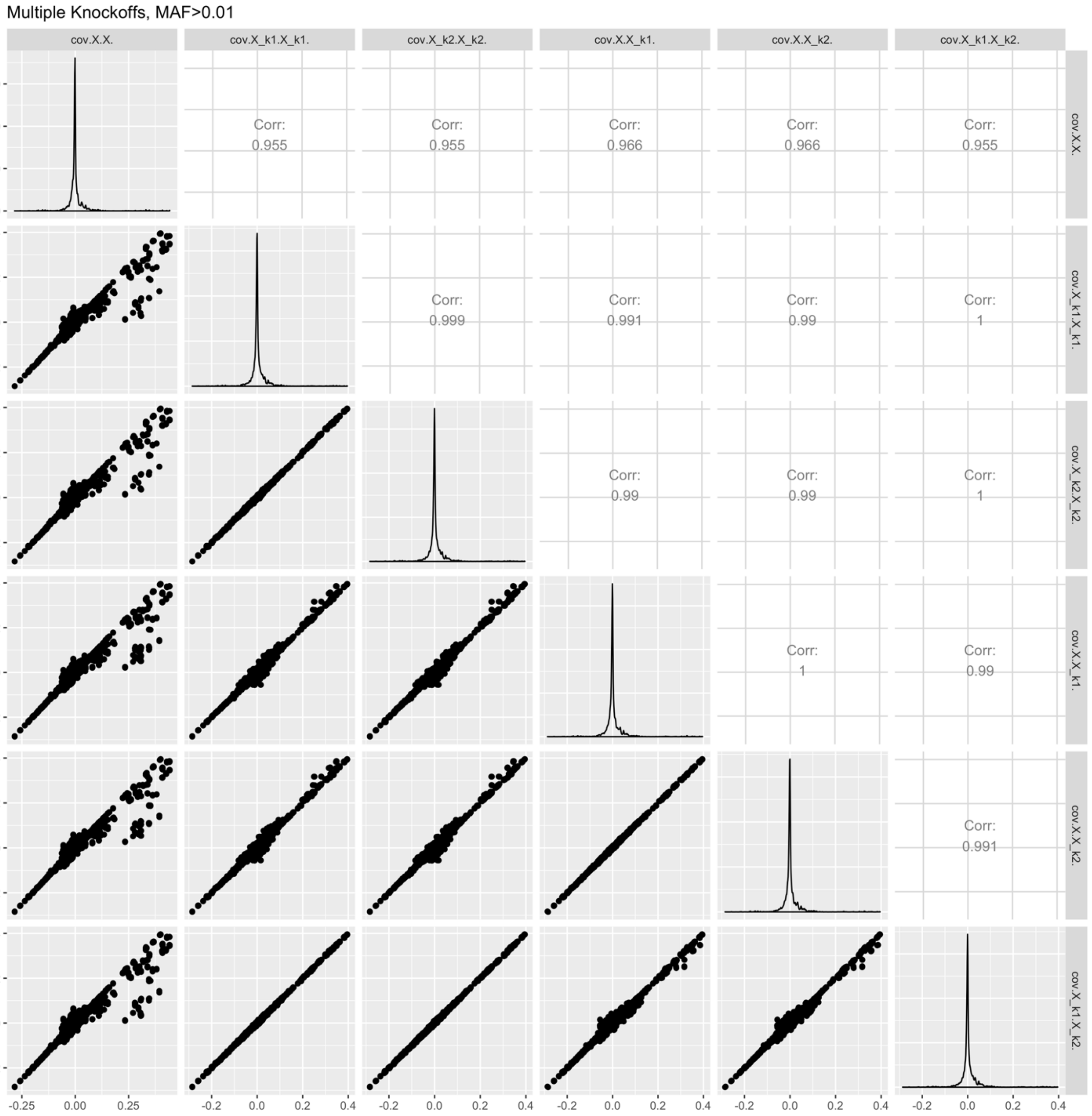

**Figure S4: *KnockoffScreen* application to the COPDGene study in TOPMed to identify variants associated with the $FEV_1$/FVC ratio in Non Hispanic White (NHW).** The top-left panel presents the Manhattan plot of p-values from the conventional association testing with Bonferroni adjustment ($p < 0.05$/number of tested windows) for FWER control. The bottom-left panel presents the Manhattan plot of *KnockoffScreen* with target FDR at 0.1. The right panel presents a heatmap that shows stratified p-values of all loci passing the FDR=0.1 threshold, and the corresponding Q-values that already incorporate correction for multiple testing. The loci are shown in descending order of the knockoff statistics. For each locus, the p-values of the top associated single variant and/or window are shown indicating whether the signal comes from a single variant, a combined effect of common variants or a combined effect of rare variants. The names of those genes previously implicated by GWAS studies are shown in bold (names were just used to label the region and may not represent causative gene in the region).

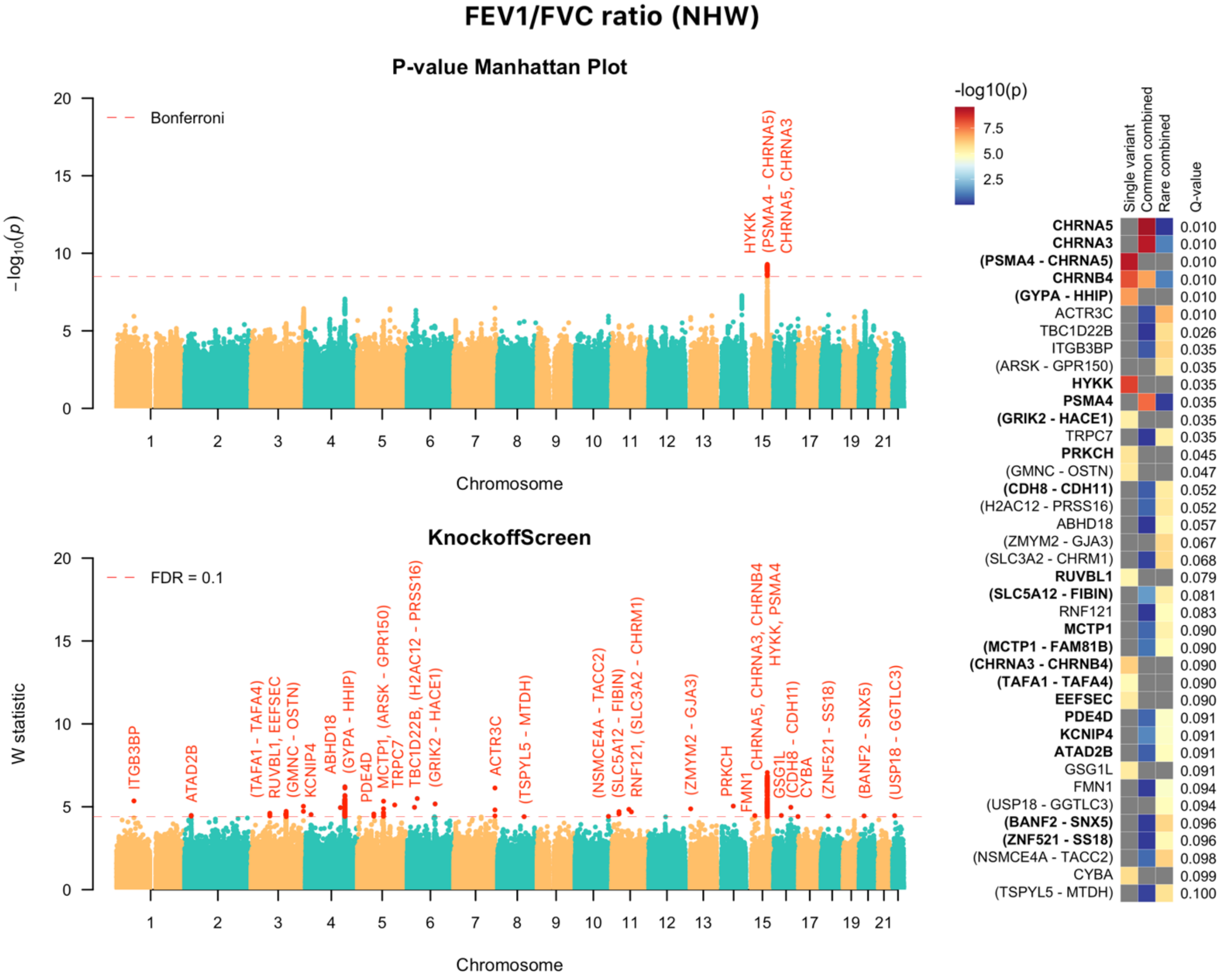

**Figure S5:** QQ plots for all tests, common variants tests, and rare variant tests included in the *KnockoffScreen* procedure for all datasets used in the analyses.

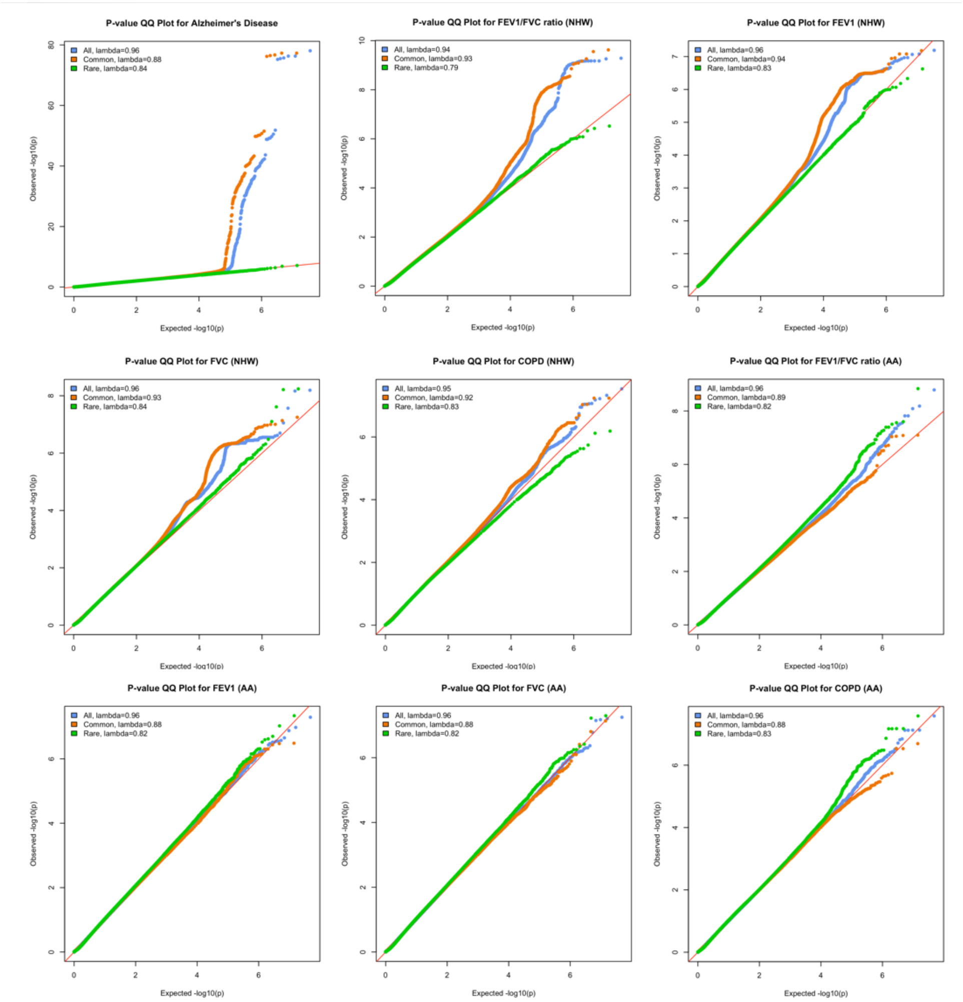

**Figure S6: The analysis of the ADSP and TOPMed data with the Benjamini–Hochberg procedure for FDR control.** The left panel presents the Manhattan plot of adjusted p-values (Q-values; truncated at $10^{-10}$ for clear visualization) from the conventional association testing with the Benjamini–Hochberg adjustment for FDR control. The right panel presents a heatmap that shows stratified p-values (truncated at $10^{-10}$ for clear visualization) of all loci passing the FDR=0.1 threshold, and the corresponding adjusted p-values that already incorporate correction for multiple testing. For each locus, the adjusted p-values of the top associated single variant and/or window are shown indicating whether the signal comes from a single variant, a combined effect of common variants or a combined effect of rare variants. The names of those genes previously implicated by GWAS studies are shown in bold (names were just used to label the region and may not represent causative gene in the region).

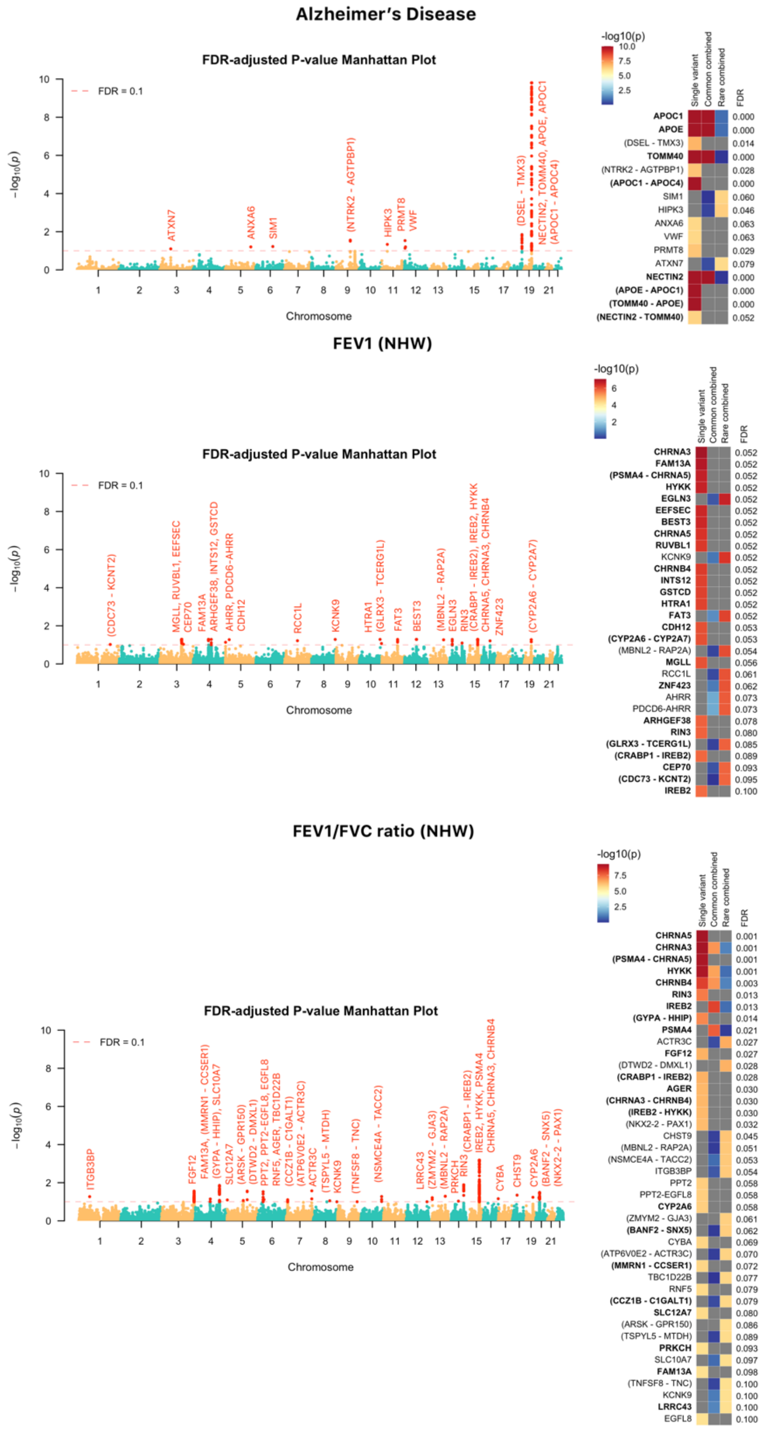

**Figure S7: Comparison with HMM (S=12) stratified by minor allele frequency.** We generated 10,000 individuals with genetic data for a 200 kb region containing 1000 genetic variants, simulated using a coalescent model (COSI). We compared the proposed algorithm to HMM with number of states S=12 and evaluated whether the second order (correlation between each pair of genetic variants) is exchangeable. Each dot presents one variant/window. The left panels evaluate how the correlation structure of knockoffs is similar to that of the original variants; the right panels evaluate how the knockoffs preserve the correlation structure when one swaps a variant with its synthetic counterpart.

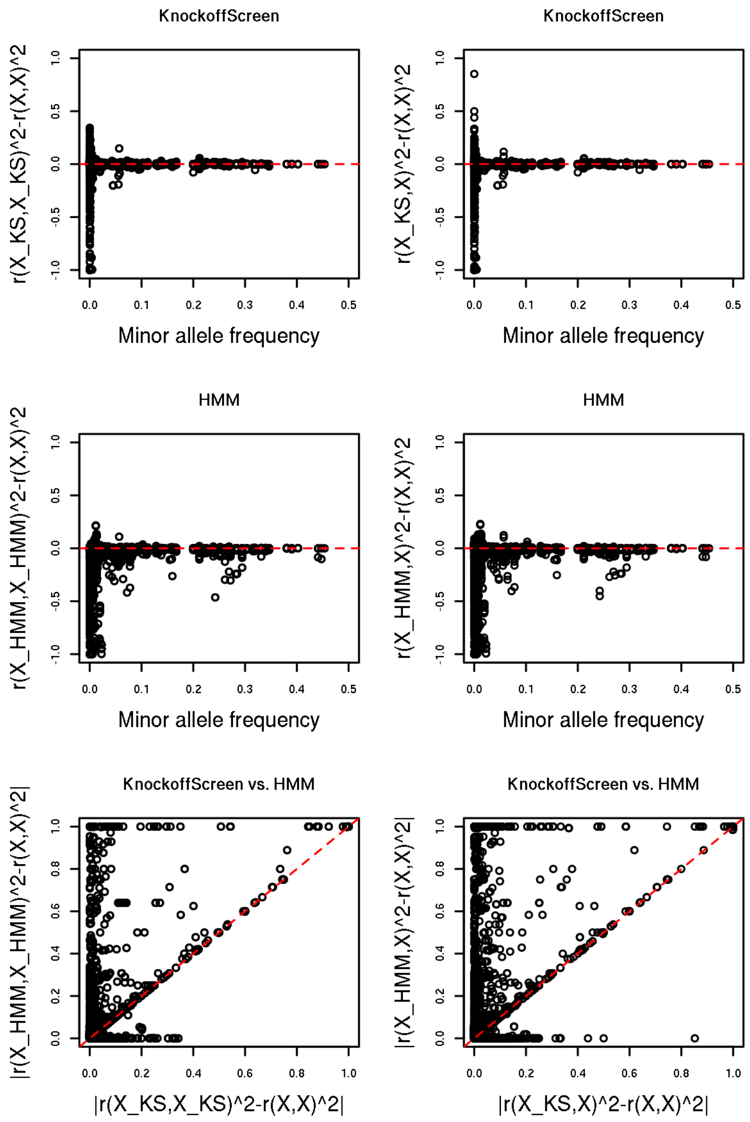

**Figure S8: Comparison with HMM (S=50) stratified by minor allele frequency.** We generated 10,000 individuals with genetic data for a 200 kb region containing 1000 genetic variants, simulated using a coalescent model (COSI). We compared the proposed algorithm to HMM with number of states S=50 and evaluated whether the second order (correlation between each pair of genetic variants) is exchangeable. Each dot presents one variant/window. The left panels evaluate how the correlation structure of knockoffs is similar to that of the original variants; the right panels evaluate how the knockoffs preserve the correlation structure when one swaps a variant with its synthetic counterpart.

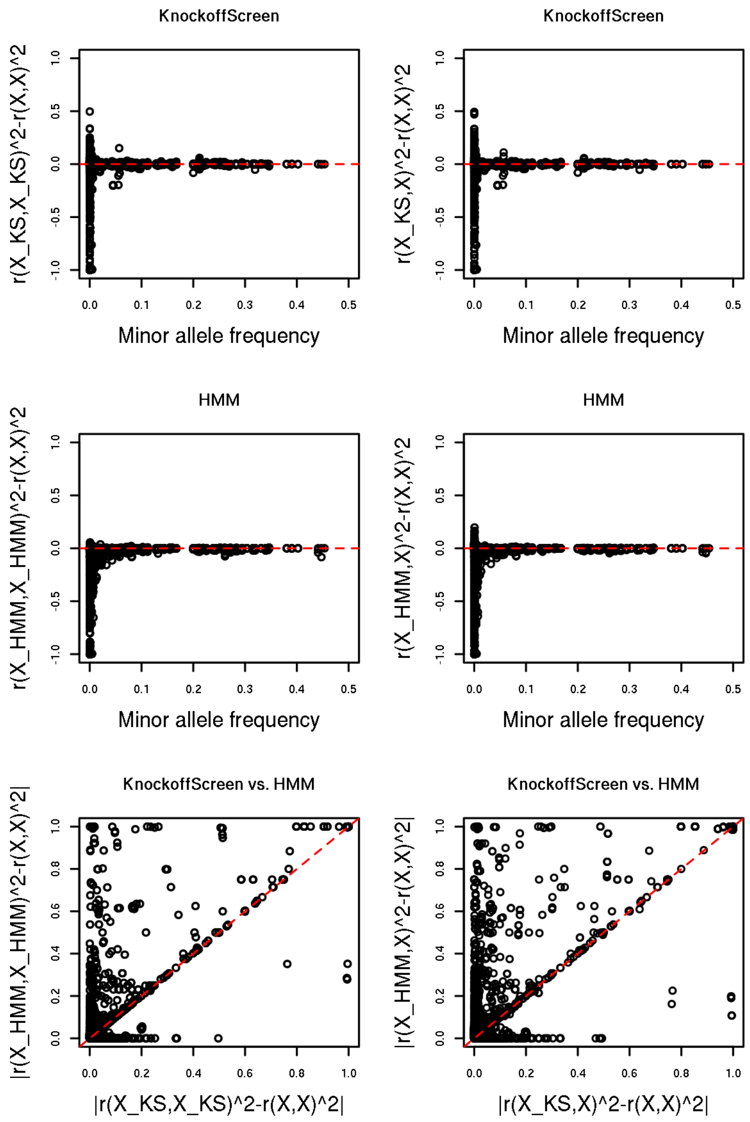
